# Human dorsal root ganglia after plexus injury: either preservation or loss of the multicellular unit

**DOI:** 10.1101/2023.02.06.526934

**Authors:** Annemarie Schulte, Johannes Degenbeck, Annemarie Aue, Magnus Schindehütte, Felicitas Schlott, Max Schneider, Camelia Maria Monoranu, Michael Bohnert, Mirko Pham, Gregor Antoniadis, Robert Blum, Heike L Rittner

## Abstract

**Objective:** Plexus injury results in lifelong suffering of flaccid paralysis, sensory loss, and intractable pain. For this clinical problem, regenerative medicine concepts, such as cell replacement for restoring dorsal root ganglion (DRG) function, set high expectations. However, it is completely unclear which DRG cell types are affected by plexus injury.

**Methods:** We investigated the cellular composition of human DRG in a clinically characterized cohort of patients with plexus injury. Avulsed DRG of 13 patients were collected during reconstructive nerve surgery. Then, we analyzed the cellular composition of the DRG with a human-adapted objective deep learning-based analysis of large-scale microscopy images.

**Results:** Surprisingly, in about half of the patients, the injury-affected DRG no longer contained DRG cells. The complete entity of neurons, satellite glial cells, and microglia was lost and replaced by mesodermal/connective tissue. In the other half of patients, the cellular entity of the DRG was well preserved. We found no loss of neurons, no gliosis, and macrophages close to single sensory neuron/satellite glial cell entities. Patients with ‘neuronal preservation’ had less pain than patients with ‘neuronal loss’.

**Interpretation:** The findings classify plexus injury patients in two categories: type I (neuronal preservation) and type II (neuronal loss). We call for early, post-accidental interventions to protect the entire DRG and improved MRI diagnostics to detect ‘neuronal loss’. Regenerative medicine to restore DRG function will need at least two translational directions: reafferentation of existing DRG units for type I injuries; or replacement of the entire DRG structure for type II patients.

## Introduction

Brachial plexus injury mainly occurs after traffic accidents and is associated with profound functional impairments. Typically, during the accident, ventral and dorsal roots are damaged at the dorsal root entry zone. In upper supraclavicular brachial plexus injury, 69 %-84 % of patients with total palsy report pain.^1, 2^ This often results in lifelong disability of patients in the middle of their life.^3^ Surgical interventions include neurolysis, nerve grafting, nerve transfer, and muscle or tendon transfer to restore motor function.^4, 5^ However, sensory function and pain relief are barely improved by the surgery and mean pain decreases only by 1.2 points on the numerical analog scale.^3^ Several concepts might explain increased pain after accidental nerve injury: Discussed are, amongst others, lack of sensory input and ectopic firing in the dorsal horn, increased activity of regenerating axons and collaterals, maladaptive peripheral nociceptor input, as well as central sensitization.^6^

To restore sensory functions, repair of different motor-, sensory and nociceptive structures of the nervous system could be helpful.^7^ Thus, neural replacement strategies set high expectations.^8^ However, up to date, the cellular changes after peripheral nerve injury in humans are not known. Without this knowledge, it is virtually impossible to design new treatments including regenerative medicine, cell-targeted immunotherapy, or pathfinding strategies.^9, 10^

Very few studies have analyzed plexus injury; so, most of the evidence has been extrapolated from firstly more peripheral nerve lesions and secondly almost exclusively from rodents. These injuries lead to adaptions in the DRG cell composition and cellular plasticity in the whole DRG.^11-14^ Loss of sensory neurons seems to depend on the severity of injury.^14^ Satellite glial cells (SGC), a cell type that forms a functional entity with the sensory neuron soma, react within hours after nerve injury and contribute to pain by enhancing neuronal excitability.^15^ Typical plasticity responses of SGCs, e.g. after ventral root avulsion or spared nerve injury,^11, 16^ include an increased expression of the glial fibrillary acidic protein (GFAP) or apolipoprotein J (APOJ).^13, 15, 17^ Furthermore, macrophages become activated, invade, or proliferate locally in the DRG close to the neuron-SGC entity.^16, 18, 19^ However, human DRG are quite distinct from rodents’,^20-22^ and it is currently barely investigated how the cellular composition of human DRG is affected by different forms of nerve injury.

Here, we clinically phenotyped a cohort of patients with brachial plexus injury. Avulsed DRG were collected during nerve reconstruction surgery and analyzed by immunohistochemistry, large-scale tile microscopy, human DRG-adapted deep learning-based algorithms,^23^ and new concepts for quantitative assessment of the different cell types close to the neuron-satellite-glia unit.^11^ Our data document that patients suffering from plexus injury can be categorized into two groups: patients with total loss or patients with complete preservation of the multicellular DRG entity.

## Materials and methods

### Patient recruitment

The study was approved by the local ethical committee (plexus injury patients: 32/19, cases from forensic autopsies: 205/20) and registered at the Germany clinical trials registry (DRKS00017266). The cohort comprised 13 patients of both sexes after brachial plexus injury, mostly due to a motorcycle accident. All patients endured multiple injuries, including avulsion of the dorsal and ventral root. Study participants gave written informed consent. For a 17-year-old patient, a parent gave consent. Three to eight months after the accident, avulsed cervical DRG were collected during reconstructive surgery. The avulsed dorsal root and the corresponding DRG tissue was located, dissected, and immediately processed.

Human control DRG were obtained from seven study subjects, 3-6 days post-mortem, during routine forensic autopsy with potential injury of the central nervous system (Institute of Forensic Medicine, University of Würzburg, Germany). One of the seven cases had to be excluded from the image analysis due to post-mortem tissue degradation. In another sample, the conglomerate of neurons in the extracted tissue was too small for systematic analysis.

### Clinical characterization and patient reported outcomes

Preoperatively, patients underwent an MRI of the plexus and answered questionnaires on patient-reported outcomes. Additionally, clinical information on age, gender, laboratory results, and patient history, including a chart review, was obtained. The following patient-reported outcomes were used in the respective German versions^24^: (1.) The graded chronic pain scale (GCPS, Von Korff Score, grade 0-IV) and scores for pain intensity (numeric rating scale, NRS).^25^ (2.) The disabilities of arm, shoulder, and hand were assessed by the DASH questionnaire. ^26^ (3.) Neuropathic pain characteristics were reported using the neuropathic pain symptom inventory (NPSI, range: 0–1).^27^ Patients were also requested to draw the localization of their pain. (4.) Depression was assessed using the Beck Depression Inventory II (BDI II, range: 0–63). A value of 0–13 was characterized as minimal, 14–19 as mild, 20-28 as medium, and 29-63 as severe depression.^28^. (5.) The State-Trait Anxiety Inventory (STAI-T, STAI-S, range 20-80) examined anxiety as a state or trait characteristic.^29^ (6.) The pain catastrophizing scale (PCS) was used with 0 being the lowest and 52 the highest pain catastrophizing.^30^

### Magnetic resonance neurography of patients after plexus injury before surgery

Patients were examined at the Department of Neuroradiology of the University Hospital Würzburg, between 06/2020 and 03/2022. MR neurography examinations were carried out on a 3 Tesla unit (MAGNETOM Skyra, Siemens Healthineers). The same examination protocol was applied to all patients and was specifically developed for posttraumatic imaging of the brachial plexus from the spinal nerve root level to the distal levels in the axilla. Part of the extensive clinical protocol was the SPACE (3D turbo spin echo with variable flip angle) STIR (short tau inversion recovery) sequence with TR/TE 2500/208 ms, TI 210 ms, parallel imaging (GRAPPA 3, reference lines PE 24), slice thickness 1.0 mm, number of slices 104, FOV 305 × 305 mm^2^, acquisition matrix 305 × 305, pixel spacing 0.95 × 0.95 mm^2^ and an acquisition time of 6:19 min. The sequence was acquired using a custom-designed coil dedicated to high-resolution imaging of the supra-, retro- and infra-clavicular brachial plexus (Variety 16-Channel Multipurpose Coil, NORAS MRI product). This two-element 2 × 8-channel surface coil was positioned on the corresponding body surface on the affected side. Multi-planar image reconstruction was performed with isotropic voxel size.

### Assessment of the extent of brachial plexus injury

On a first level of analysis, we counted all dorsal root avulsions, regardless of whether they were stated as complete, partial, or suspected in the presurgical MRI and doctor’s reports. In a second analysis, we added the number of segments that were affected by any kind of lesion, e.g., peripheral nerve or spinal cord injuries, or bone fractures. Areas of pain or sensitivity alterations were obtained from the patients’ and doctors’ reports and assigned to corresponding dermatomes. Sensitivity changes included hyperalgesia, allodynia, dysesthesia, a positive Tinel’s sign, hypoesthesia, or a complete loss of sensitivity.

### Tissue preparation

Immediately after tissue harvest, human DRG were transferred into ice-cold 4 % paraformaldehyde/PBS solution. DRG were transported on ice and were fixed overnight at 4°C. The next day, the tissue was washed three times for 30 min in phosphate-buffered saline (PBS) and stored overnight in 30 % sucrose (PBS-buffered) for cryoprotection (4°C). Afterward, the DRG was cut longitudinally into two pieces. Each half was embedded in O.C.T. compound (Tissue-Tek) in a plastic embedding mold. Freezing was performed in 2-Methylbutane, pre-chilled with liquid nitrogen. The tissue was stored in a -80°C freezer. DRG were serially sectioned at 20 μm and mounted on twelve SuperFrost Plus slides (Menzel). Each slide contained a representative collection of 3–4 slices from different areas of the corresponding DRG. After sectioning, the slides were immediately used for immunofluorescence staining.

### Hematoxylin and eosin staining (H&E)

Frozen sections were swayed for a short period of time in a 4 % formaldehyde solution and afterwards in water. Then, the slides were put for 1–2 min into Mayer’s hematoxylin solution. After washing in water, the slides were swayed shortly in a 1 % eosin solution and were again washed in water. Then, the sections were swayed first in ethanol and afterward in xylol. Labeled sections were finally embedded in a xylene-based mounting medium (Pertex).

### Immunofluorescence

Slides were incubated for 15 min in quenching solution (7.5 g/l glycine in aqua dest., pH 7.4). Labelling was performed with standard methods. In brief: blocking solution was 10 % donkey or horse serum, 0.3 % Triton X100, 0.1 % Tween 20 in PBS.). Primary and secondary antibodies were used in blocking solution (**Supplementary Table 1**). Cell nuclei were stained with DAPI (0.1 mg/l in PBS; 7 min). Slides were washed in PBS, shortly desalted in water, and finally embedded in Aqua Polymount (Polysciences).

**Table 1.** Clinical characteristics of study subjects.

| Parameter | Patients with plexus injury (n = 13) |
| --- | --- |
| Age (years) | 34.7 ± 13.8 |
| Sex | f: 2, m: 11 |
| Type of accident | motorcycle: 8, other: 5 |
| Time since injury (months) | 4.7 ± 1.3 |
| **Number of dorsal roots affected** |  |
| Affected by Dorsal Root Avulsion | 2.8 ± 1.2 |
| Affected by all lesions | 4.6 ± 0.9 |
| **Pain** |  |
| Current pain (numeric rating scale, NRS 0-10) | 4.7 ± 2.2 |
| Mean pain (numeric rating scale, NRS 0-10) | 5.0 ± 2.3 |
| Maximum pain (numeric rating scale, NRS 0-10) | 7.6 ± 1.8 |
| Graded chronic pain scale (GCPS, I-IV) | 2.8 ± 1.2 |
| **Disability and neuropathic pain** |  |
| Disability arm, shoulder, and hand (DASH): total symptom score | 51.2 ± 15.4 |
| Disability arm, shoulder, and hand (DASH): work score | 80.0 ± 22.2 |
| Neuropathic pain symptom inventory (NPSI, 0-1) | 0.3 ± 0.2 |
| **Psychiatric comorbidities** |  |
| State-Trait Anxiety Inventory: State (STAI-S, 20-80) | 45.5 ± 13.3 |
| State-Trait Anxiety Inventory: Trait (STAI-T, 20-80) | 41.7 ± 15.7 |
| Beck depression inventory 2 (BDI II, 0-63) | 14.8 ± 13.9 |
| Pain catastrophizing scale (PCS, 0-52) | 22.5 ± 12.1 |
| **Treatment (number of patients)** |  |
| Non-opioid analgesics | 11 |
| Opioids | 4 |
| Anticonvulsants | 9 |
| Antidepressants | 7 |
| Cannabinoids | 1 |
| No analgesics | 0 |
| **Control group (n = 5)** |  |
| Age (years) | 68.0 ± 23.5 |
| Sex | f: 2, m: 3 |
| Cause of death | accident: 2, accident or cardiovascular disease: 1, suicide: 1, cerebral venous sinus thrombosis: 1 |
| Interval before autopsy (IBA, days) | 3.8 ± 1.3 |
| Interval before autopsy without cooling | 1 h of IBA: 2, 2 h of IBA: 2, n. a.: 1 |

GCPS: grade I-IV; I, low disability-low intensity; II, low disability-high intensity; III, high disability-moderately limiting; IV, high disability-severely limiting. NPSI: no cut-off. STAI: STAI-S ≤ 39 defined as normal. BDI II: 0-13 minimal, 14-19 mild, 20-28 moderate, 29-63 severe depression. PCS: 0-29 low, 21-37 moderate, 38-52 high catastrophizing tendency. DASH: no cut-off. Treatment refers to medication prescribed at hospital discharge after reconstructive surgery, multiple medications possible. All data are mean ± standard deviation.

### Microscopy and quantitative image analysis

Tile scan microscopy was performed with an Axio Imager 2 microscope (Zeiss), a Plan Apochromat 20× (N.A. 0.8) objective, and an Axiocam 506 camera (monochromatic, 14-bit). Pixel resolution was 0.454 μm. Objective bioimage analysis was performed using *deepflash2*^31^ as described recently,^11, 23^ with some modifications. Image feature annotation was performed on 10 exemplary sections marked by neurofilament (NF) by three experts using the QuPath software (Bankhead et al. 2017, RRID:SCR_018257). The ground truth of the annotations of the three experts was estimated with the simultaneous truth and performance level estimation (STAPLE) method (**Supplementary Table 2**). The *deepflash2* software created a deep learning model from these annotations.^31^ Training of a new DL model ensemble (three models) was performed with eight of the ten exemplary images (**Supplementary Table 3**). Two images were used to test the model. Annotation overlap and the model’s performance were evaluated with the Dice score, defined as: 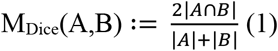; where A and B are two sets of pixels. Then, the model ensemble was applied to predict the NF-positive neurons for all images.

IBA1-positive macrophages with thin cell extensions were challenging for manual annotations. However, due to the high quality of the labels with a high signal-to-noise ratio, masks could be created with a thresholding method. First, NF-positive segmented regions were excluded from the IBA1 images to remove falsely/lipofuscin-stained parts. Then, scikit-image filters were used for background reduction and sharpening (unsharp_mask function). Calculation of an “optimal” threshold was computed with the Otsu method. Pixels with fluorescence intensity above the Otsu Threshold were considered IBA1-positive. Similarly, but without excluding neuronal regions, the masks for FABP7-positive SGC were created. Thresholding was sufficient because FABP7 almost exclusively stained SGC with a high signal-to-noise ratio. Even though the filters accounted for a lot of the viability, the DRG images of one control needed to be excluded from the subsequent analysis because the thresholding method did not work adequately for images with a low signal-to-noise ratio.

Finally, the predicted areas of neurons, SGC, and macrophages were used for image feature quantification. For this, python scripts were created https://github.com/AmSchulte/DRGhuman. The neuron-near area (NNA) was calculated through binary dilation (scikit-image morphology) of the NF segmentation. As another reference area, a convex hull was drawn to define the neuron polygon area (NPA). For SGC quantification, the area, and the proximity to neurons (percentage of neurons in proximity to FABP7-positive SGC), were used. Both the NF mask and the SGC mask were dilated by one pixel and if the overlap between a dilated neuron and the surrounding FABP7 mask was bigger than 0, the neuron was counted to be in proximity to a FABP7-positive SGC. The IBA1 area was quantified inside the NNA and the intensity of the APOJ signal was determined within the FABP7-positive SGC normalized to the rest of the APOJ intensity in the NNA. Calculated image parameters were averaged for each DRG.

### Statistical analysis

The statistical analyses were performed using the GraphPad Prism software, version 8.4.3. The clinical and histological data were first tested for normal distribution by the Shapiro-Wilk test. Neuronal soma sizes, due to many individual data points, were tested for normal distribution with the D’Agostino-Pearson test (Omnibus K2). Normally distributed data were tested with the F-test for homoscedasticity. Normally distributed, homoscedastic datasets were tested for significance by the unpaired, two-tailed t-test. Normally distributed datasets with unequal variance were tested for significance by the unpaired, two-tailed t-test with Welch’s correction. Significance tests for non-normally distributed data were performed by the two-tailed Mann-Whitney test. The distribution underlying the Bouhassira cluster data among the groups ‘neuronal loss’ and ‘neuronal preservation’ was tested by the two-sided Fisher’s exact test. Correlation analyses were performed using the Spearman test. A p-value < 0.05 was considered statistically significant. In boxplots, single data points are presented. The boxes extend from the 25^th^ to the 75^th^ percentile and the lines within the boxes show the median. The whiskers extend from the smallest to the highest value.

### Data availability

Images and corresponding segmentation as well as the DL-models are available at Zenodo.^32^ The python scripts of the analysis are openly available on GitHub: https://github.com/AmSchulte/DRGhuman.

### Ethics statement

The authors declare the compliance with ethical standards including informed consent and adherence to good clinical practice.

## Results

### Patients with plexus injury: clinical data of the cohort

The plexus injury group was comprised of 13 mostly male patients – on average five months after brachial plexus injury, typically due to a motorcycle accident (**Table 1**). Patients suffered overall from complete avulsion of two to three dorsal roots. Usually, five segments were affected by the trauma.

Patients described presurgically moderate mean pain, high maximum pain, and moderate neuropathic pain quality (NPSI) (**Table 1**). Half of the patients reported a higher level of state anxiety – four of these even moderate to severe depressive symptoms (**Table 1**). Pain catastrophizing (PCS) was on average below the threshold. The DASH score revealed that all patients were highly impaired in their upper extremity activity – in daily life and even more at work.

All patients received pain medication. The most frequently prescribed drug class was non-opioid analgesics, followed by anticonvulsants and antidepressants. Four patients received opioids and one cannabinoids (**Table 1**). The control cohort data were acquired from five cases scheduled for forensic autopsy (**Table 1**). The control group was 27.8 years older than the patient cohort.

### Either ‘neuronal loss’ or ‘neuronal preservation’ in patients with plexus injury

Human DRG showed fiber-rich regions interrupted by neuron-rich regions (**Fig. 1**). Neuron-rich areas were predominantly localized in the periphery of DRG sections. A thick protective connective tissue layer surrounded DRG and was between neuronal somata. All neuronal somata contained a nucleus with a prominent nucleolus and were surrounded by SGC. Surprisingly, in seven of the 13 patient tissue samples, neither neurons nor SGC were found. To be sure that this was true for the whole DRG, we stained more serial slices with H&E, but could not find any neuron/SGC entities. Only fat cells and connective tissue were found next to nerve fibers. In controls obtained from forensic autopsy, DRG tissue was preserved and comparable to that of plexus injury patients with neuronal preservation.

**Figure 1.**
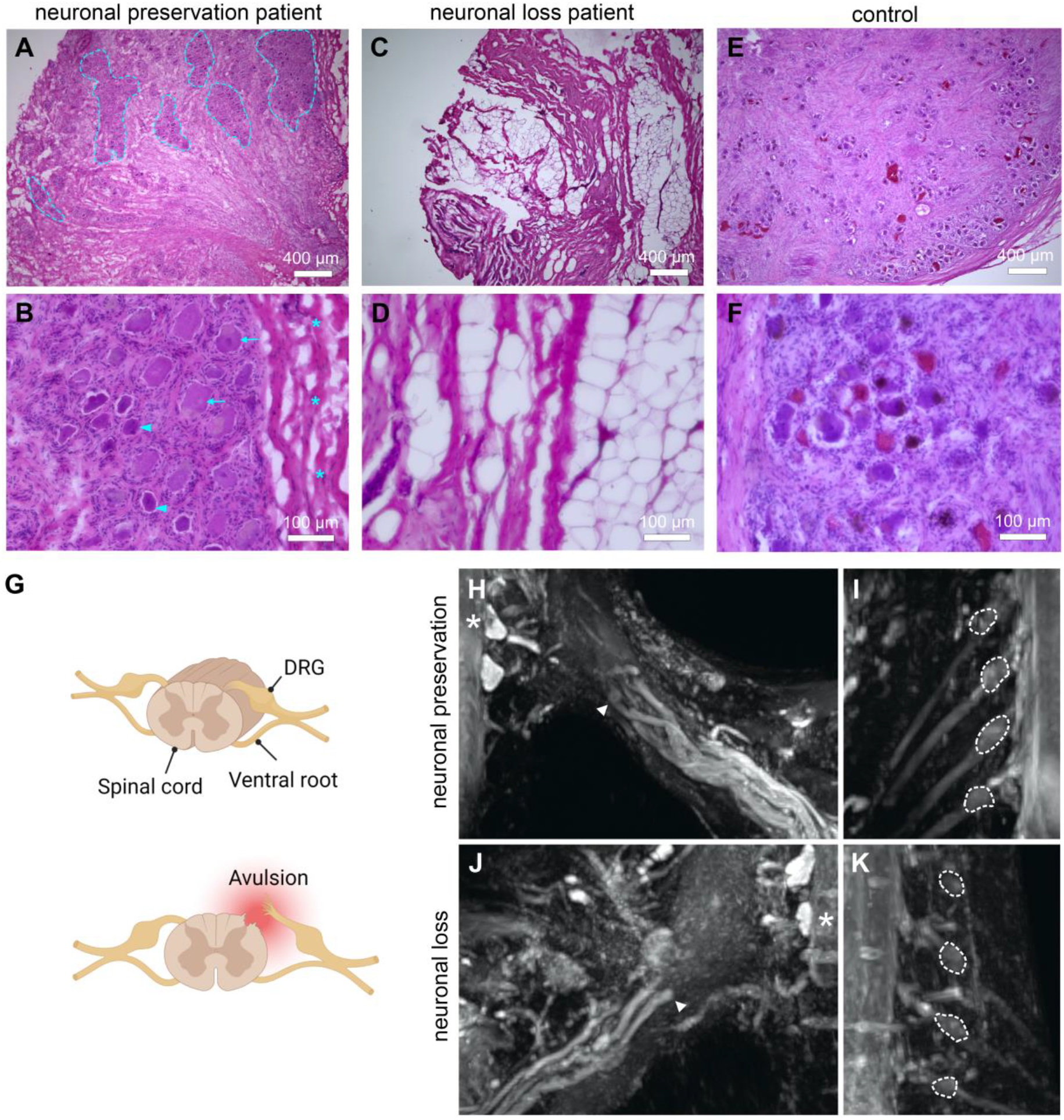
Classification into ‘neuronal preservation’ or ‘neuronal loss’. (**A-F**) Representative images of H&E-stained sections of DRG from plexus injury patients. Indicated are: neuron-rich area (outlined in A), small sensory neurons (arrowheads in B), large sensory neurons (arrows in B), connective tissue (asterisks in B). (**G**) Scheme of DRG depicting a dorsal root avulsion. Created with biorender©. (**H-K**) Images resulting from magnetic resonance neurography of the brachial plexus in two different dorsal root avulsion patient groups. In both groups, affected DRG can no longer be identified, nor their volume quantified. Indicated are nerve stumps (arrowheads in H and J) and pseudomeningoceles of C8 roots (asterisks in H and J) as well as unaffected DRG at C5-C8 of the contralateral side (dotted line in I and K).

Presurgical magnetic resonance neurography did not allow reliable delineation of the affected DRG from injured adjacent neural structures in either ‘neuronal loss’ or ‘neuronal preservation’ patients posttraumatically. In Fig. 1h, a ‘neuronal preservation’-patient with an intradural avulsion of the left C8 and T1 roots is shown. Here, the retracted nerve stumps form a neuroma and did not allow delineation of the affected DRG due to post-traumatic fibrous scarring and adhesions. Fig. 1j is representative of a ‘neuronal loss’-patient. An intradural avulsion of the right C7 and C8 root is visible, retracted nerve stumps formed a neuroma and we could not delineate the affected DRG. On the contralateral sides, the unaffected DRG, segments C5-C8, could be properly identified.

In the clinical examination, areas of pain and loss of sensation did not match injured segments (**Fig. 2**): Most patients felt pain and sensitivity alterations in dermatomes of both injured and non-injured segments. Even when all kinds of lesions – avulsions of the corresponding dorsal or ventral roots, more peripheral nerve lesions, lesions of the spinal cord or bone fractures – were considered, pain and sensitivity alterations were reported in dermatomes outside of the injured segments (5 out of 13 for pain and 3 out of 13 for sensation). This supports the concept of somatosensory plastic changes after plexus injury.

**Figure 2.**
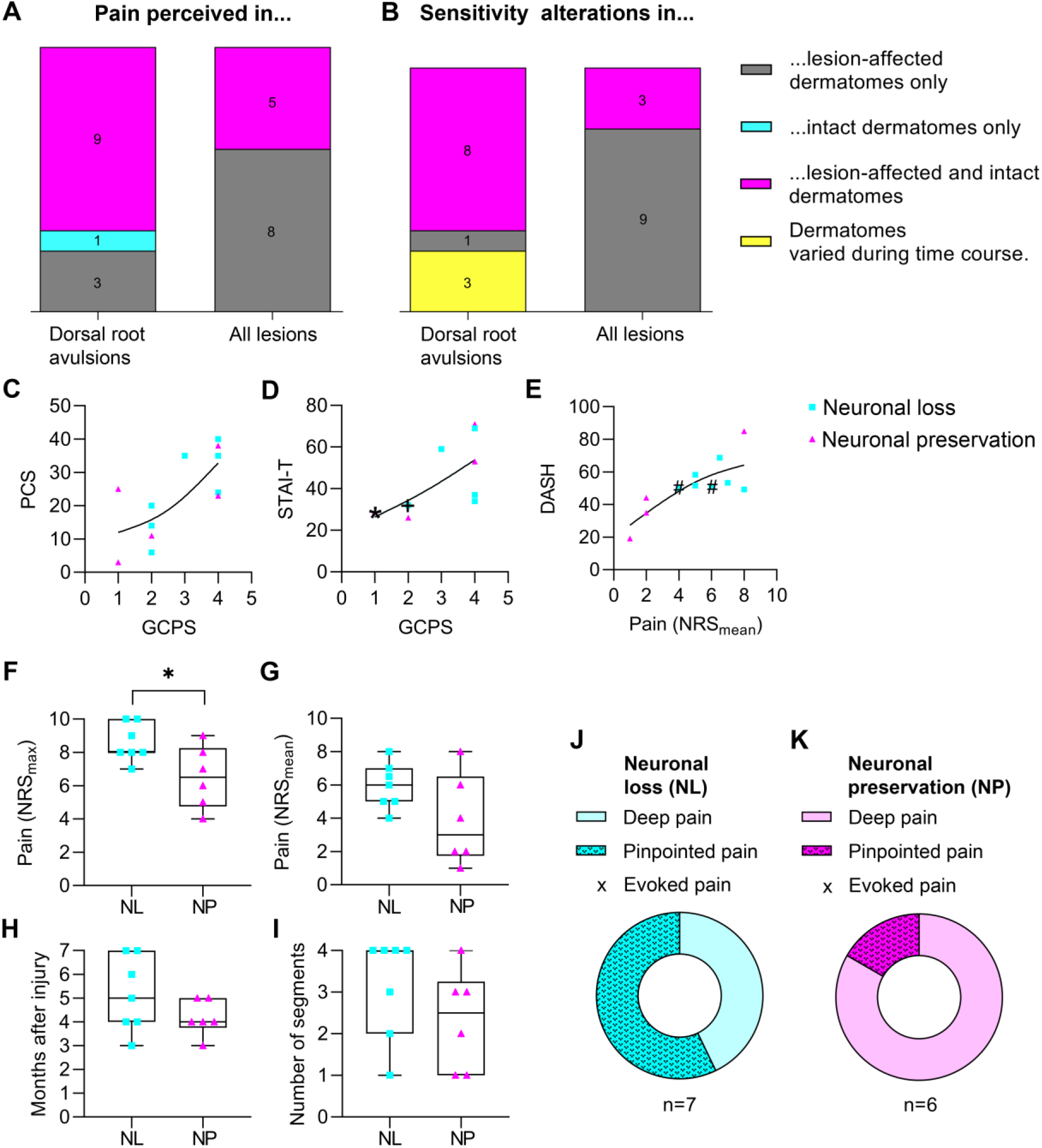
Extension of clinical symptoms beyond the injury and increased severity in patient with ‘neuronal loss’. (**A**) Areas of reported pain matched to dermatomes. (**B**) Areas of sensitivity alterations, e. g. allodynia or dysesthesia, or numbness, matched to dermatomes. Sensitivity alteration data of one patient was not available. Patients were assigned to four categories. In grey: Patients with symptoms only in dermatomes supplied by corresponding avulsed dorsal roots. In cyan: patients with symptoms only in dermatomes supplied by intact roots. In magenta: patients with symptoms in both types of dermatomes supplied by intact or avulsed roots. In yellow: patients who reported different sensitivity changes in the time course between accident and surgery. Left columns: dermatomes normally supplied by avulsed dorsal roots, right columns: dermatomes potentially affected by any kind of lesion. (**C-E**) Correlation of pain with psychiatric comorbidities in patients after plexus injury. (**C**) Pain Catastrophizing Scale (PCS) in relation to von Korff’s Graded Chronic Pain Scale (GCPS, Spearman test, r=0.6751, p=0.0197). (**D**) Trait-anxiety of the State-Trait-Anxiety Inventory (STAI-T) with the GCPS (Spearman test, r=0.8405, p=0.0013). Overlay of single patient data: **\*** marks two ‘neuronal preservation’ patients; **+** marks three ‘neuronal loss’ patients. (**E**) Disability of the Arm, Shoulder, and Hand score (DASH disability/ symptom score) in comparison to mean pain on the Numeric Rating Scale (NRS) during the last four weeks before surgery (Spearman test, r=0.6981, p=0.0098). Patients with type II ‘neuronal loss’ are depicted in cyan, the type I ‘neuronal preservation’ group in magenta. Trends are shown as black line. ^**#**^ marks overlay of single data points: ‘neuronal loss’ with a ‘neuronal preservation’ patient. (**F-K**) Comparison of clinical data of patients with type I ‘neuronal preservation’ or type II ‘neuronal loss’. (**F**) Maximal pain on the numeric rating scale (NRS) during the last four weeks before surgery (unpaired, two-tailed t-test, *p = 0.0317). (**G**) Mean pain according to NRS (unpaired, two-tailed t-test, p = 0.0990, not significant). (**H**) Time between injury and reconstructive surgery in months (unpaired, two-tailed t-test, p = 0.1939, not significant). (**I**) Number of dorsal roots affected by dorsal root avulsion (two-tailed Mann-Whitney test, p = 0.2541, not significant). (**J, K**) Neuropathic pain phenotype according to Bouhassira clusters. No patient reported evoked pain. Two-sided Fisher’s exact test, p = 0.2657, not significant. Number of patients: ‘neuronal loss’ group: n = 7; ‘neuronal preservation’ group: n = 6.

We next classified the patients into group ‘type I - patients with neuronal preservation’ and ‘type II – patients with neuronal loss’. Unrelated to the classification, psychiatric comorbidities influenced pain and functional impairments (**Fig. 2**): Trait-anxiety and catastrophizing correlated significantly with the GCPS. The highest activity impairment was reported by patients with symptoms of depression and anxiety. Mean pain intensity was associated with functional upper extremity impairment evaluated in the DASH score.

However, type I patients showed a trend to more pain, injury, and more analgesic treatment (combination of anticonvulsants and antidepressants, **Fig. 2, Supplementary Table 4**). The majority of the type II patients with ‘neuronal loss’ perceived pain as ‘deep’ feeling pressuring and squeezing as measured by the clusters of the NPSI,^27^ whereas most ‘neuronal preservation’ patients perceived pain as ‘pinpointed’ reporting paresthesia. Of note, none of patients was in the ‘evoked pain’ cluster – experiencing pain provoked by brushing, pressure, or cold.

### Cellular plasticity and satellite glia cell marker distribution in human DRG

Next, we validated plasticity markers by triple color immunofluorescence in confocal microscopy. *Neurofilament* (NF) and *microtubule-associated protein 2* (MAP2) were reliable markers for neurons and labeled sensory neuron somata of all different sizes (**Fig. 3**). Intracellular lipofuscin aggregates with substantial autofluorescence were observed in all samples. Almost all sensory neuron somata were surrounded by *fatty acid binding protein 7* (FABP7)-positive SGC, whereas *glutamine synthetase* (GS) and *glial acidic fibrillary protein* (GFAP) signals were seen far more sparsely in both patient and control DRG (**Fig. 4**). Anti-APOJ (*clusterin*) immunoreactivity also formed ring-like features surrounding NF-positive neuronal somata and was largely co-localized with the SGC marker FABP7. Macrophages labeled by spindle-shaped IBA1 (*allograft inflammatory factor 1* or *ionized calcium-binding adapter molecule 1*) were present in the DRG around NF- and MAP2-positive neuronal somata. Some macrophages were also seen in the surrounding connective tissue.

**Figure 3.**
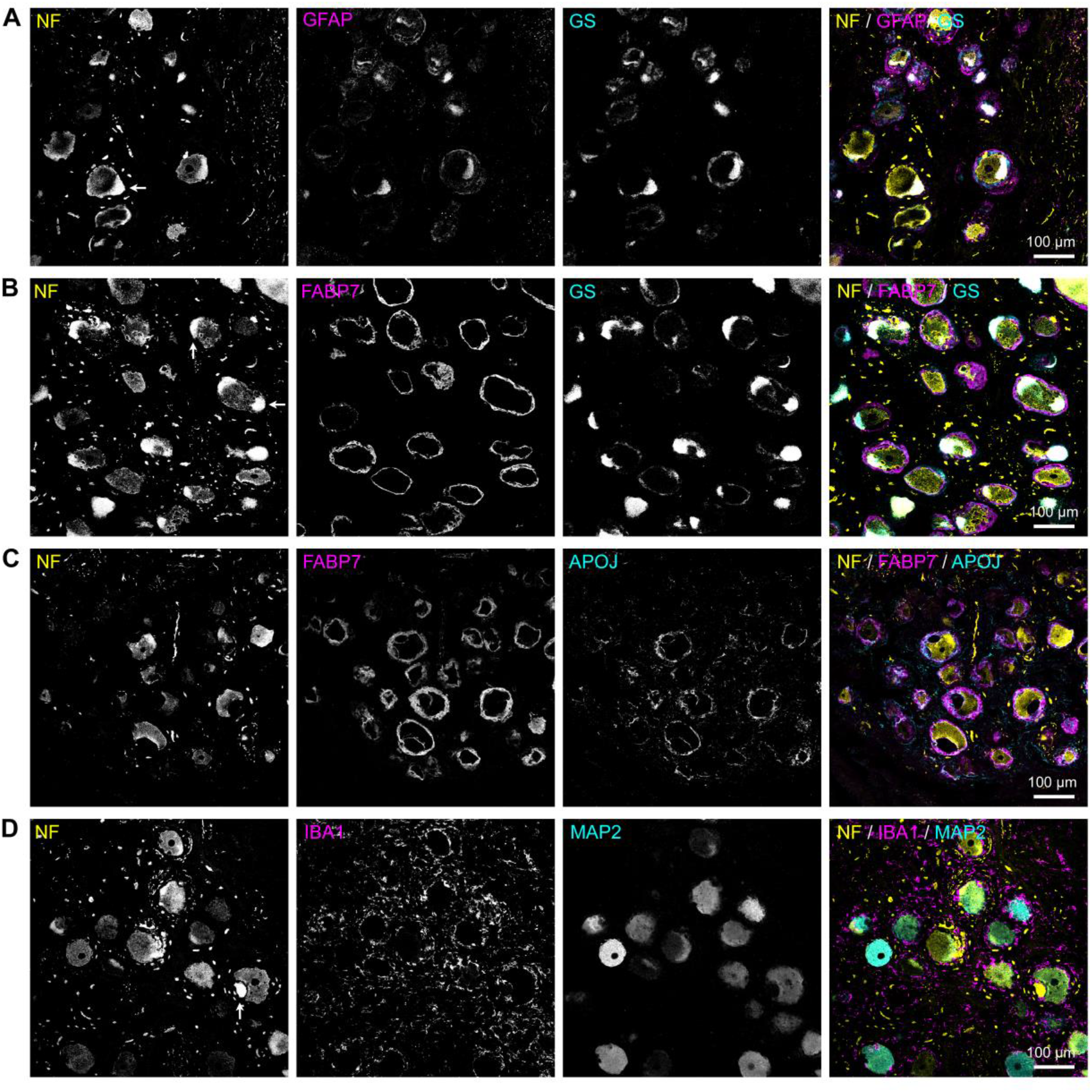
Immunofluorescence labeling of human DRG. (**A-D**) Representative confocal images (maximum intensity projection) show DRG labels from plexus injury patients. Neurons were labeled with indicated markers. NF (neurofilament, yellow, in A-D) and MAP2 (microtubule-associated protein 2, cyan, in D). Satellite glial cells were marked using GFAP (glial fibrillary acidic protein, magenta, in A), GS (glutamine synthetase, cyan, in A), FABP7 (fatty acid binding protein 7, magenta, in B, C) and APOJ (clusterin, cyan, in C). Macrophages are labeled using IBA1 (ionized calcium binding adaptor molecule 1, in D). Lipofuscin-mediated autofluorescence was observed in all tissue samples (see arrows). Scale bar: 100 μm.

**Figure 4.**
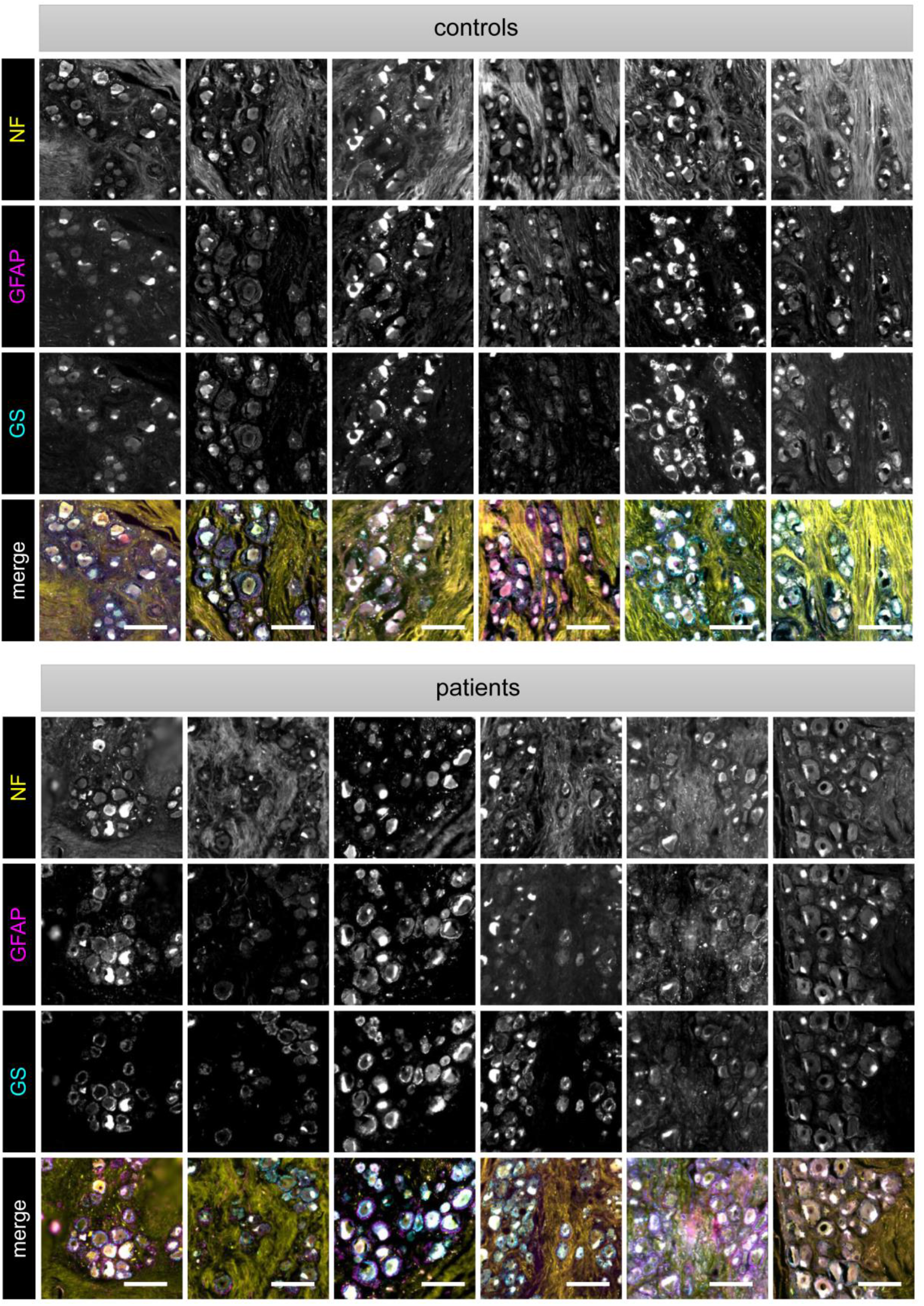
Large heterogeneity of immunoreactivity to GFAP and GS in human DRG slices. Sections of the six controls and six plexus injury patients from the ‘neuronal preservation’ group stained for glial fibrillary acidic protein (GFAP) and glutamine synthetase (GS). Scale bar: 200 μm.

### Comparable cellular composition in the ‘neuronal preservation’ group

Tissue sections from different patients showed a high heterogeneity (**Fig. 4**). To objectively analyze microscopy images showing the cellular composition of the human DRG (**Fig. 5**), we adapted a recently introduced deep learning (DL)-based analysis strategy.^11^ First, DL models needed to be trained to allow computational segmentation of the neurons in the human DRG. For this, three experts annotated NF-positive neurons in representative human DRG images. Subsequently, a computational consensus information of all experts (ground truth) was computed and used to train deep learning (DL)-model ensembles. This strategy has been confirmed to increase the objectivity and validity of bioimage analysis.^23, 33^ Expert annotations overlapped highly with the ground truth estimation (**Supplementary Table 2**). DL models, validated with different ground truth image data sets, reached a mean *dice score* of 0.875 (**Supplementary Table 3**), thus confirming reliable sensory neuron segmentation in the human DRG. Testing of the DL-model ensemble was performed on annotated images that were previously not seen by the DL-model. This control test showed that the trained DL-model predicted segmentations of NF-positive neurons in the human DRG with human expert-like performance (**Fig. 5**).

**Figure 5.**
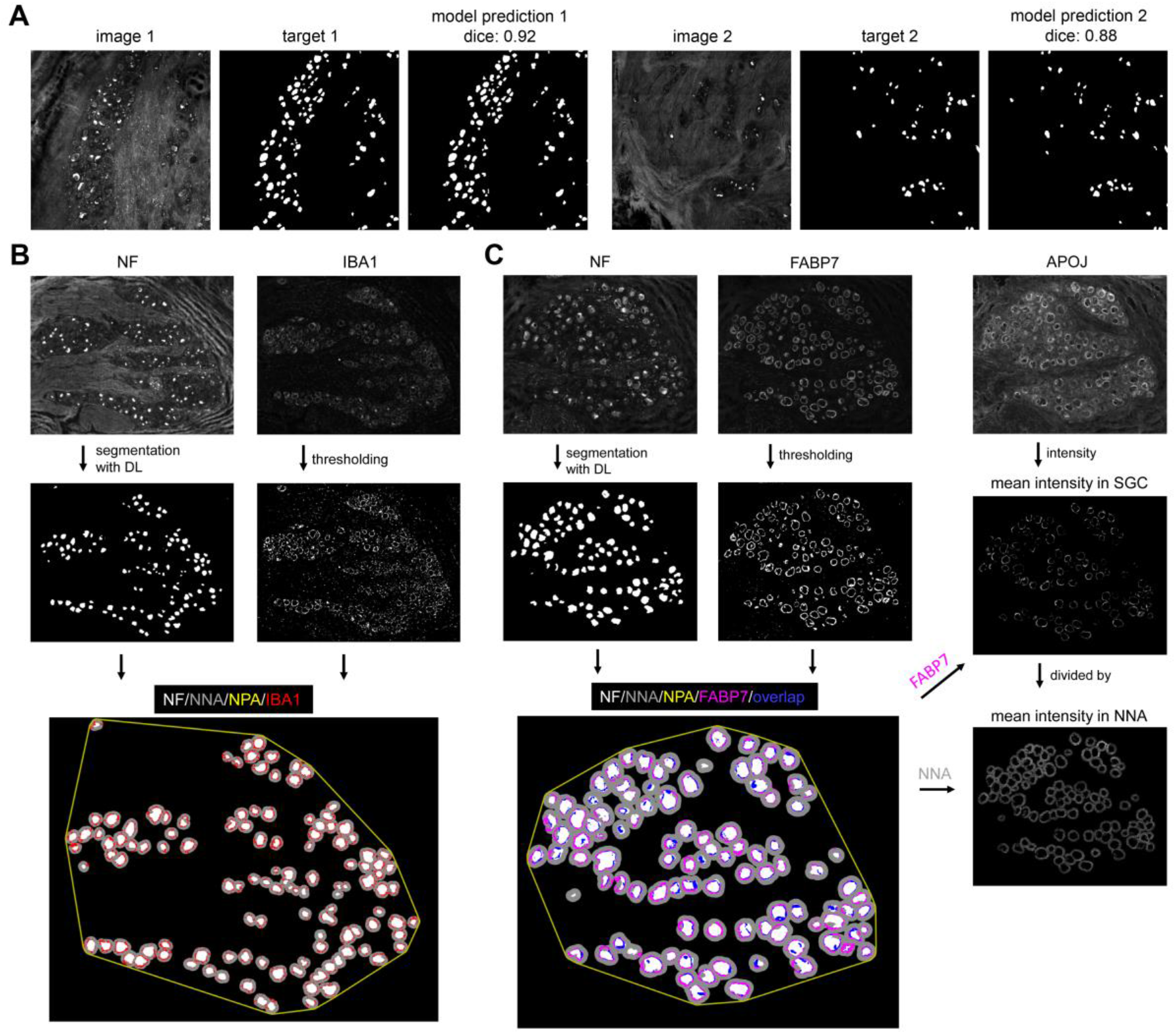
Analysis procedure for neurofilament-segmented DRG sections. (**A**) Performance of the deep learning model ensemble in predicting neurofilament (NF) labels on two exemplary test images.Shown are NF images, their expert-annotated segmentation target (ground truth) and the corresponding model prediction. Similarity of target and model prediction was evaluated with the dice score. (**B**) NF in IBA1-labeled sections was segmented with the DL model ensemble. IBA1 was segmented with a thresholding method. Neurons (NF, white) were quantified in the neuronal polygon area (NPA, yellow), which is computed with a convex polygon around segmented neurons. The IBA1-positive area (red) was quantified inside and in relation to the neuron near area (NNA, grey). (**C**) Sections labeled for satellite glial cells (SGC) were analyzed for the FABP7-positive area and APOJ signal intensity. NF was segmented with the DL model ensemble and FABP7 with a thresholding method. NF-positive neurons (white) were quantified in relation to the neuronal polygon area (NPA, yellow). For FABP7, the signal-positive area per neuron near area (NNA) and the percentage of neurons in proximity to FABP7-positive SGC (overlap in blue), was determined. The APOJ intensity was calculated by dividing the mean intensity in FABP7-positive SGC by the mean intensity in the neuron near area (NNA).

Neuronal somata of control DRG were slightly smaller than those DRG of patients with ‘neuronal preservation’ but the number of neurons did not differ (**Fig. 6**). Almost all (95 %) of the DRG neurons were in proximity to SGC (FABP7-positive SGC), showing that the entity of sensory neurons and their SGC was preserved. Furthermore, similar numbers of FABP7-positive cells occupied the neuron near area, indicating the absence of gliosis. No upregulation of the injury-related factor *clusterin* (APOJ) in SGC was observed. A trend for more macrophage abundance close to neurons was found in the patients. The data show that the multicellular composition of the DRG, consisting of sensory neurons of different sizes, their SGC ring, and neuron-near macrophages, was preserved in patients after plexus injury.

**Figure 6.**
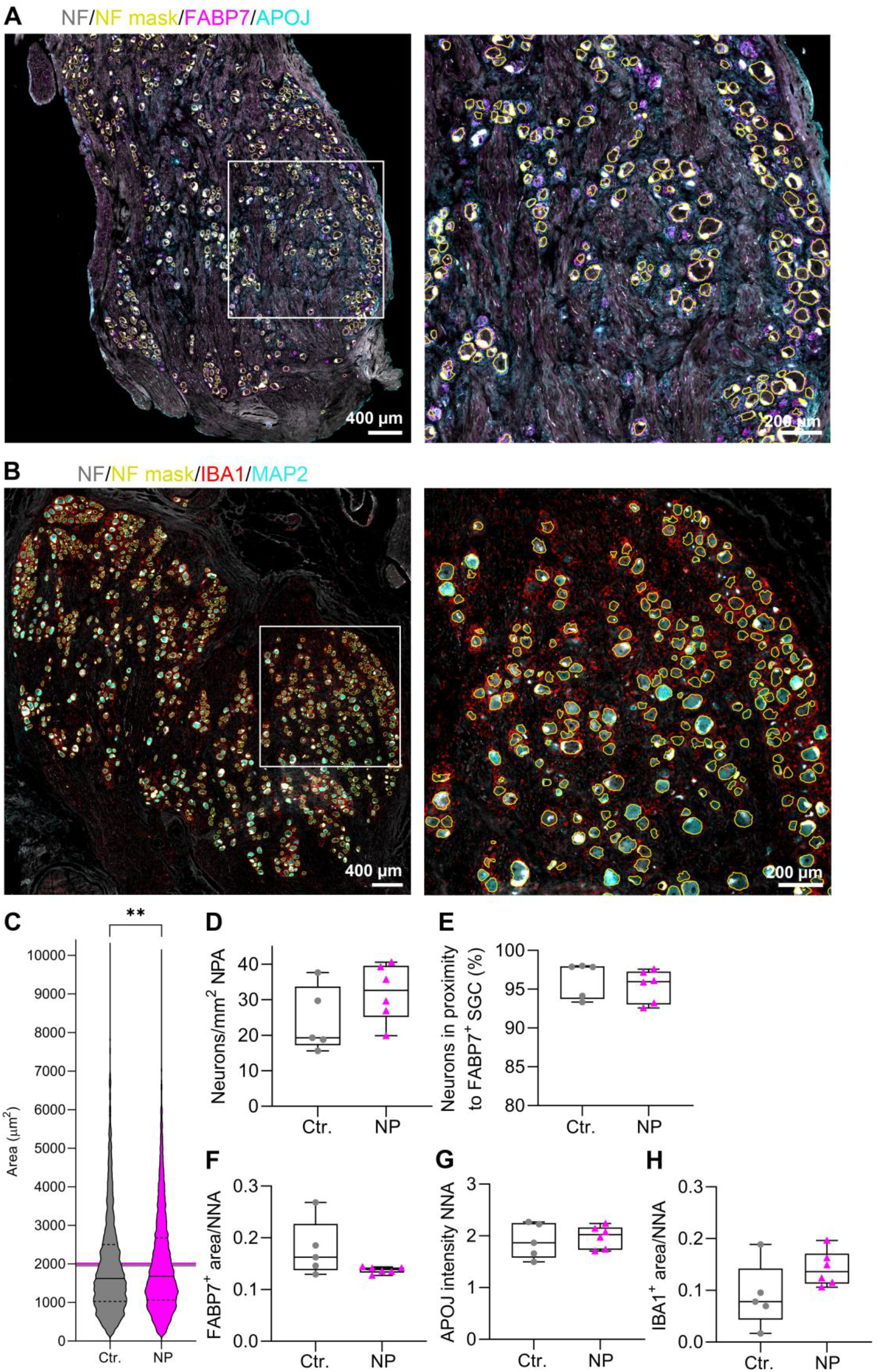
Similar cellular composition of DRG from patients with ‘neuronal preservation’ vs. control. Deep learning-based immunohistochemical analysis comparing control (Ctr., grey, circles, n = 5) versus ‘neuronal preservation’ DRG (NP, magenta, triangles, n = 6). (**A**) Representative images of a patient DRG section with neurons (NF, grey) and SGC (FABP7, margenta; APOJ cyan). NF-positive neurons were segmented with a deep learning model (NF mask, yellow). (**B**) Labels for neurons (NF, grey; NF-mask, yellow; MAP2, cyan) and macrophages (IBA1, red). (**C**) Violin plot showing the soma size of neurons. Black lines: median area of neuronal somata, dashed lines: 25th and 75th percentile, lines in grey and magenta: mean area of control and ‘neuronal preservation’ DRG (two-tailed Mann-Whitney test, **p = 0.0050). (**D**) Number of neurons per mm^2^ in the neuron-rich area (computed as neuronal polygon area, NPA) (unpaired, two-tailed t-test, p = 0.1689, not significant). (**E**) Percentage of neurons in proximity to FABP7-positive satellite glial cells (SGC, two-tailed Mann-Whitney test, p = 0.2468, not significant). (**F**) Neuron near area (NNA) occupied by FABP7-positive cells (unpaired, two-tailed t-test with Welch’s correction, p = 0.1698, not significant). (**G**) Intensity of the anti-APOJ signal in SGC normalized to the intensity of the APOJ signal in the neuron near area (NNA, unpaired, two-tailed t-test, p = 0.6681, not significant). (**H**) Neuron near area (NNA) occupied by IBA1 positive macrophages (unpaired, two-tailed t-test, p = 0.1111, not significant).

## Discussion

Here, we surprisingly found that patients with plexus injury belong to two dichotomous groups: patients with plexus injury type I with ‘neuronal preservation’ and type II with ‘neuronal loss’ in their extracted DRG. Contrary to our hypothesis, in DRG with ‘neuronal preservation’, we observed neither less neurons, nor significant macrophage invasion, nor upregulation of GFAP in SGC, nor gliosis. This injury at the border between the PNS and the CNS seems to evoke an ‘all-or-none’ response. We do not know the reason why some of the patients lost their DRG tissue, but these patients had a tendency towards more widespread injury. Thus, it is possible that more injury resulted in hematoma formation, disruption of blood supply, more inflammation and scarring, mitochondrial collapse, and anoxic death. Brachial plexus injuries are rather common, have a bad prognosis and are a tremendous economic burden, as the injury happens very often in young people.^34-36^ Our observation has huge implications for therapy; hence, we introduced a novel cellular classification of preganglionic plexus injury: type I with ‘neuronal preservation’ and type II with ‘neuronal loss’.

The clinical presentation of our cohort was representative: all patients scheduled for surgery had pain with some neuropathic symptoms and were largely impaired in daily life.^1, 3^ Among the typical disease burden coming from flaccid paralysis, numbness, and intractable pain, a quarter of our patients reported symptoms of depression and increased anxiety levels – as shown in other cohorts.^37^ Accordingly, we think that about half of the corresponding patients worldwide are type II patients and suffer from a loss of avulsed DRG tissue.

Our patients reported squeezing and pressure pain, comparable with ‘deep pain’ in the NPSI clusters.^27^ Pain was not restricted to the corresponding dermatomes of the injured segments. This could be due to collateral damage, root overstretching, or plastic changes due to aberrant innervation from collateral or regenerating nerves. Damaged neurons are capable of spontaneous or ectopic activity causing spontaneous or shooting pain.^38^ Input from nociceptors is necessary because the application of local anesthetics to adjacent roots relieves pain independent of the individual somatosensory phenotype or signs of central sensitization.^39^ Disruption of the pain pathway might also spread to other ipsilateral DRG or may lead to central sensitization and/or ectopic activity originating in the dorsal horn.

Pain is the major limitation for return to work and quality of life after plexus injury. Thus, treatment of neuropathic pain remains the major clinical problem here. Most of our patients received anticonvulsants and non-opioids, although, for neuropathic pain, antidepressants have a better efficacy (lower number needed to treat) and non-opioids are not effective.^40^ Conceptually, treatment aiming at a restoration of sensory and nociceptive pathways might help here – as it is observed that neonates with plexus injury have an excellent sensory recovery and never develop pain.^41^ This argues that an intact somatosensory system is necessary for pain relief,^41^ thus making neural replacement strategies after sensory function loss conceptually attractive.

In preclinical models, the ‘neuronal loss’ phenotype has, to our knowledge, never been observed. SGC are activated in response to peripheral postganglionic and preganglionic injury as well as dorsal or ventral root avulsion.^16, 42, 43^ In human tissue, we did not detect any signs of gliosis using typical SGC markers like GS, and FABP7. GS/GLUL is found in 50 % of rodent and 10 % of human SGC – both on mRNA and protein.^44^ GS showed a very low expression in only a few SGC in plexus injury patients and in healthy human DRG – so it does not seem to be a good marker in humans. Across species, 88-96 % of neurons are surrounded by FABP7-positive SGC. This is in line with our data showing 95 % SGC-neuron entities (neurons in proximity to FABP7-positive SGC) in the type I ‘neuronal preservation’ patient group and comparable numbers in the forensic medicine controls. We saw no change in APOJ expression in SGC around neurons in the human DRG. No GFAP, the known SGC activation marker,^17^ was described so far in uninjured human SGC.^44^ Interestingly, we see some GFAP abundance in all samples - independent of injury. Additionally, we found local macrophages close to the neuron/SGC entity, in line with others.^45^ Thus, the cellular DRG unit with neurons, SGC, and local microglia is preserved in some patients, whereas it completely vanishes in others.

Our study has several limitations. In our á priori planning, we did not expect to find patients with a complete loss of DRG neurons. Therefore, the two patient groups are rather small, hence, the clinical data have low statistical power. This could not be avoided due to a rare disease and limitations in the ethical approval, because DRG after plexus injury are typically not removed during nerve reconstruction. Our patient sample was therefore restricted to a small number of patients to allow hypothesis-generating research. Secondly, DRG after plexus injury were compared with post-mortal DRG (3-6 d after death). These control samples were not easy to match with the experimental DRG samples. Moreover, human DRG tissue contains a substantial amount of connective tissue and is quite variable, making it difficult to keep the image exposure time constant across individual sections. To counteract this, we have adapted an – at least on the image level – objective analysis procedure, which is based on feature extraction in ambiguous bioimage data.^11, 31^ Human DRG tissue samples are so valuable that the bioimage data, the deep learning models, and the annotated images are helpful research tools in our open science community. They can be shared and reused, thus supporting research on human DRG on the way to new therapies.

In summary and importantly, our discovery asks for early intervention strategies to prevent DRG tissue loss after injury. To get visual control over this clinical process, we need to consider early and better MR-imaging applications for injury diagnosis. Future MRI methods should allow improved quantification of the DRG’s cell body-rich area to stratify the neuronal status non-invasively, at best over the course of the disease, to better understand when and why neuron/SGC entities are lost.^46^ For patients with type II ‘neuronal loss’, tissue replacement may be a strategy. However, for restoring DRG neurons, the complete neuron-SGC entity would need to be replaced. Here, the use of neural crest-derived sensory progenitors could be considered as a regenerative medicine concept, as these cells can give rise to both SGC as well as all types of DRG neurons.

## Abbreviations

APOJ: apolipoprotein J/clusterin
DASH: Disability arm, shoulder, and hand
DL: deep learning
DRG: dorsal root ganglia
FABP7: fatty acid binding protein 7
GFAP: glial fibrillary acidic protein
GS: glutamine synthetase
IBA1: ionized calcium-binding adapter molecule
MAP2: microtubule-associated protein 2
NF: neurofilament
NL: neuronal loss
NNA: neuron near area
NP: neuronal preservation
NPA: neuronal polygon area
NPSI: Neuropathic pain symptom inventory
NRS: numeric rating scale
ROI: regions of interest
SGC: satellite glial cells

## Acknowledgements

We thank all patients and their physicians for participating in this study. We acknowledge Olivia Rudtke for her technical help with the DRG analysis. We would like to thank Brigitta Grolik as well as the staff of the Peripheral Nerve Surgery Unit, University of Ulm. We are grateful to Heiko Besenfelder and Max Perschneck from the Institute of Forensic Medicine, University of Würzburg, for their help with the human DRG biopsy.

AS was funded by the Evangelisches Studienwerk Villigst. JD was funded by the Graduate School of Life Sciences, University of Würzburg. RB, and HLR were funded by the Interdisciplinary Center for Clinical Research Würzburg (IZKF) project N-D-368. MP, RB, and HLR were supported by the German Research Foundation, project ID: 426503586, KFO5001 ResolvePAIN.

## Author contributions

Conceptualization: AA, AS, JD, CM, HLR, RB; Methodology: AA, AS, JD, RB, HLR; Investigation: AA, AS, JD, RB, FS, MS, MB, GA, HLR; Software: AS; Formal analysis: AA, AS, JD; Funding acquisition: HR, RB, AS, MP, JD; Supervision: HLR, RB, CM, MP; Project administration: HLR, RB; Writing – original draft: AA, AS, JD; Writing – review & editing: HLR, RB

### Conflicts of Interest

The authors declare no conflict of interest.

## Notes

### Competing Interest Statement

The authors have declared no competing interest.

