## Supplementary Table 4 for "Human dorsal root ganglia after plexus injury: either preservation or loss of the multicellular unit"

**Supplementary Table 4 Clinical characteristics of patients with 'neuronal loss' and 'neuronal preservation'**

| | **Type I neuronal preservation (n = 6)** | **Type II neuronal loss (n = 7)** |
|---|---|---|
| Age (years) | 37.5 ± 14.3 | 32.3 ± 14.0 |
| Sex | f: 1, m: 5 | f: 1, m: 6 |
| **Number of dorsal roots affected** | | |
| Affected by all lesions | 4.5 ± 1.2 | 4.7 ± 0.5 |
| **Pain** | | |
| Current pain (Numeric rating scale 0-10) | 3.7 ± 2.6 | 5.5 ± 1.6 |
| Graded Chronic Pain Scale (GCPS) | 2.4 ± 1.5 | 3.0 ± 1.0 |
| **Disability and neuropathic pain** | | |
| Disability arm, shoulder, and hand (DASH): total symptom score | 47.2 ± 21.9 | 54.6 ± 6.9 |
| Disability arm, shoulder, and hand (DASH): work score | 68.8 ± 28.4 | 87.5 ± 15.3 |
| Neuropathic pain symptom inventory (NPSI, 0-1) | 0.3 ± 0.3 | 0.3 ± 0.1 |
| **Psychiatric comorbidities** | | |
| State-Trait Anxiety Inventory: State (STAI-S, 20-80) | 47.2 ± 15.7 | 44.1 ± 12.0 |
| State-Trait Anxiety Inventory: Trait (STAI-T, 20-80) | 41.2 ± 17.7 | 42.1 ± 15.3 |
| Beck depression inventory 2 (BDI, 20-63) | 15.2 ± 14.8 | 14.3 ± 14.5 |
| Pain catastrophizing scale (PCS, 0-52) | 19.7 ± 12.1 | 24.9 ± 12.5 |
| **Treatment** | | |
| Number of classes | 1.8 ± 0.8 | 3.0 ± 1.3 |
| Non-opioid analgesics | 4 | 7 |
| Opioids | 1 | 3 |
| Anticonvulsants | 4 | 5 |
| Antidepressants | 2 | 5 |
| Cannabinoids | 0 | 1 |
| Anticonvulsants and antidepressants | 1 | 4 |
| No analgesics | 0 | 0 |

Cut-offs see Table 2. Treatment refers to medication prescribed at hospital discharge after reconstructive surgery, multiple medications possible. All $p > 0.05$. Unpaired, two-tailed t-test or two-tailed Mann-Whitney test. All data are mean ± standard deviation.
