## Supplementary Table 1 for "Human dorsal root ganglia after plexus injury: either preservation or loss of the multicellular unit"

**Supplementary Table 1 Primary and secondary antibodies**

| Antibodies | Company | Cat. No. | RRID | Host | Concentra-tion | Dilution |
|---|---|---|---|---|---|---|
| **Primary** | | | | | | |
| Anti-APOJ | Abcam | ab39991 | AB_722878 | Goat | 0.5 mg/ml | 1:50 |
| Anti-NFh | Millipore | AB5539 | AB_11212161 | Chicken | 0.5 mg/ml | 1:200 |
| Anti-GS | BD Transduction | 610517 | AB_397879 | Mouse | 0.25 mg/ml | 1:200 |
| Anti-GFAP | OriGene | DP-014 | AB_1001789 | Rabbit | N.A. | 1:100 |
| Anti-FABP7 | Invitrogen | PA5-24949 | AB_2542449 | Rabbit | 0.29 mg/ml | 1:100 |
| Anti-IBA1 | Fujifilm Wako | 019-19741 | AB_839504 | Rabbit | 0.5 mg/ml | 1:100 |
| Anti-MAP2 | Sigma | M1406 | AB_477171 | Mouse | 0.5 mg/ml | 1:500 |
| **Secondary** | | | | | | |
| Anti-chicken 488 | Jackson IR | 703-545-155 | AB_2340375 | Donkey | 0.5 mg/ml | 1:500 |
| Anti-mouse Cy5 | Jackson IR | 715-175-150 | AB_2340819 | Donkey | 0.5 mg/ml | 1:500 |
| Anti-rabbit Cy3 | Jackson IR | 711-165-152 | AB_2307443 | Donkey | 0.5 mg/ml | 1:500 |
| Anti-goat 647 | Thermo Fisher | A21447 | AB_2535864 | Donkey | 0.89 mg/ml | 1:500 |
