## Supplementary Table 2 for "Human dorsal root ganglia after plexus injury: either preservation or loss of the multicellular unit"

**Supplementary Table 2: Ground truth estimation performance**

| expert | average dice score | std dice score |
|---|---|---|
| 1 | 0.945 | 0.013 |
| 2 | 0.931 | 0.015 |
| 3 | 0.856 | 0.030 |
| **mean** | **0.911** | **0.019** |
