## Supplementary Table 3 for "Human dorsal root ganglia after plexus injury: either preservation or loss of the multicellular unit"

**Supplementary Table 3: Model validation performance**

| file | model no. | dice score | uncertainty score |
|---|---|---|---|
| 0003.tif | 1 | 0.810 | 0.047 |
| 0007.tif | 1 | 0.865 | 0.042 |
| 0002.tif | 2 | 0.929 | 0.027 |
| 0009.tif | 2 | 0.930 | 0.030 |
| 0004.tif | 3 | 0.845 | 0.040 |
| 0006.tif | 3 | 0.870 | 0.039 |
